# Structural and functional microbial community response to short-term impact of switching between tillage and no-tillage at soil aggregate level

**DOI:** 10.1101/2020.08.03.234534

**Authors:** Juan Pablo Frene, Luciano Andrés Gabbarini, Luis Gabriel Wall

**Affiliations:** Laboratorio de Bioquímica y Biologia de Suelo, Centro de Bioquímica y Microbiología del Suelo, Departamento de Ciencia y Tecnología, Universidad Nacional de Quilmes, Bernal B1876BXD, Buenos Aires, Argentina.- CONICET, Buenos Aires, Argentina

## Abstract

An understanding of the distribution of soil microorganisms and enzyme activities at different soil aggregate level could help to understand the mechanisms operating behind different tillage management on soil structure and function. Our objective was to determine if the microbial community structure and soil enzymes activity (EA) associated with different aggregate fractions changed within the transition at switching between no-till and conventional tillage at 30 months after the switch of management on a base line field of 27 years long were no-till (NT) and conventional tillage (CT) were side by side compared. Part of NT plot was turned into new CT (n-CT) while part of CT plot was turned into new NT (n-NT). Aggregate fractions of 2000-250, 250-63, 63-20, 20-2 and 2-0.1μm were obtained from soil samples taken at 30 months after the switch. Specific microbial abundances, measured by qPCR, and EAs were greatest on 2-0.1μm following by 20-2and 2000-250μm. The EAs showed the highest activities in the CT (β-Glucosidase and β-D-cellobiosidase) and in the nCT (Phosphatase and N-acetyl-β-Glucosaminidase) in 2000-250μm. In contrast, in the intermediate fractions (250-63μm and 63-20μm), the highest activities were observed in NT soil. Microbial communities were significantly different among different aggregates. In the 20-2μm fraction, fungi were able to differentiate between current treatments, and bacteria and archaea showed similar trends. In 2000-250μm, the treatments were associated by their historical management, and the abundances in CT samples were superior to those of the NT. In contrast, in the fractions 250-63 and 63-20μm, the NT samples showed greater abundances to those of the CT and the new treatment samples have suffered differences from historical treatments. In conclusion, tillage systems influenced the spatial distribution of soil enzymes as well as the abundances of microbial communities in the different soil aggregate size fractions.

**Highlights:**

- Soil microbial structure and functional activity showed a heterogeneous distribution within aggregate soil fraction.
- Distribution of functions and microbial structure is shaped by tillage soil management
- The greater values of microbial abundance and soil activity appeared at smaller aggregates.
- Fungi abundance significantly enhanced under NT than the CT at 20-2 μm.

## 1. Introduction

Land-use change is one of the main driving factors affecting structure and functioning of soil (Tilman et al., 2002). Tillage systems modify and address the structure and ecological services of agricultural environments. Conventional tillage (CT) disrupts the soil and promotes the breakdown of plant residues, increases soil erosion and degradation, accelerates soil organic matter (SOM) mineralization and declines soil aggregates (Roldán et al., 2005; Yin et al., 2017). No-tillage (NT) appeared to work out negative effects produced by CT (Martinez et al., 2017; Triplett and Dick, 2008). NT, whereby seeds are planted directly into the residue of previous crops (Derpsch et al., 2010), can conserve moisture, alter soil temperature and increase SOM (Lal, 2015a). NT combined with crop rotation constitute a conservative and sustainable way of doing agriculture, and the conversion from conventional tillage to no-till has led many benefits like avoid soil degradation, increase SOM, aggregation and enhance soil health (Derpsch et al., 2010) and also show significant differences at the prokaryotic community level (Sengupta and Dick, 2015; Sengupta et al., 2020). However, the transition to a new tillage systems is not immediate and many studies had explored this changes at SOM containing (Alvarez & Steinbach, 2009; Díaz-Zorita et al., 2002; Du et al., 2014; Franzluebbers & Arshad, 1996; Martinez et al., 2020; West & Post, 2002), at microbial community’s composition analy ed by PLFA (Feng et al., 2003; Jiang et al., 2011b; Simmons & Coleman, 2008; Wortmann et al., 2008) and at functional level through the use of enzymes activities (Gabbarini, *in revision*; Pandey et al., 2014). Gabbarini et al. (*in revision*) showed a completed transition at biochemical level through enzymatic activities from one to another tillage system after 3 years. In addition, the conversion to a new treatment is expected to result in a significant restructuring of bacterial community composition and diversity in soil.

Soil structure is an important property that mediates many soil chemical and biological processes (Mikha & Rice, 2004) that has been recently recognized as the result of microbial activity (Lehmann et al., 2017; Lehmann et al, 2020). Soil aggregates, at different size level, constitute the structural blocks of bulk soil (Hemkenmeyer et al., 2015). Different aggregate size fraction have a differential mineralogy composition, and also, abundance and type of SOC, nitrogen content, water holding and O_2_ concentration (Allison & Jastrow, 2006; Beare et al., 1994; Christensen, 2001; Kandeler et al., 1999; Lagomarsino et al., 2012). Macroaggregates (>250 μm) protect plant-derived and contain fresher labile SOM and plant residues. Large (<250 μm) and small (<50 μm) microaggregates are formed by microbial induced bonding of clay particles, polyvalent metals and organometal complexes. (Lavallee et al., 2020). Chemical and physical properties are heterogeneous in the different fractions (Jillig et al., 2019).

In recent years, few studies have addressed the distribution of soil microbes and their activity across different micro-environments (Bach et al., 2018; Frene et al., 2018; Neuman et al 2013). Each aggregate fraction represents a distinctive habitat space for microbial communities (Bach et al., 2018). And this idea is reflected as each micro-environment supported a differential bacterial community composition (Ranjard et al., 2000). For instance, *Actinobacteria* showed a high abundance in macroaggregates and *Acidobacteria* were associated with microaggregates <53 μm from red sandy soils (Trivedi et al., 2017). Therefore, β-*Proteobacteria* was highest in microaggregates (avinic et al., 2012), on the contrary, α-*Proteobacteria* was highest in macroaggregates (Trivedi et al., 2015) both aggregates from clay soils. Shifts in the microbial community composition have important implications for soil functioning since different phylum would produce and contribute to different soil enzymes which are involved in the dynamics of nutrients in soil (Finn et al., 2017). Enzymes activities (EAs) also showed a heterogeneous distribution in micro-environments (Bustos & Perez-Mateos, 2000). The specific distribution of soil enzymes among the different aggregates has been interpreted depending on substrate availability and the type of association of enzymes with soil particles (Lagomarsino et al., 2012). Nevertheless this distribution of EAs among different size aggregates has been described with contrasting results. Some authors found that highest EAs in smaller aggregates (Trivedi et al., 2015), while other authors found highest activities in bigger ones (Zhang et al., 2016). Soil enzyme activities change according to the season and the nutrients availability (Bach and Hofmockel, 2015; Nicolás et al., 2012), thus any comparison about EAs distribution among different soil aggregates must be done within a particular nutrient status and sampling season.

The aim of this work was to explore tillage effects on soil microbial structure and soil physiological activity during the early transition to NT or CT at different aggregate scale fractions. We hypothesized that the change of tillage system will influence the microbial community composition and both soil EAs and soil physiological profile, differentially at the aggregate level scale. Therefore, the primary objective of this research was to evaluate the impact of new tillage systems on microbial communities and EAs at aggregate size fraction. To accomplish this goal, we fractionated soil samples in five different aggregates fractions and evaluated microbial community through molecular approaches like qPCR in concert with soil biochemical (soil enzyme (Marx et al., 2005) and physiological analysis) approaches to determine the effect of tillage systems.

## 2. Materials and Methods

### 2.1. Site and sampling

The field experiment was carried out on a loamy Typic Argiudoll at Tornquist (38° 07’ 06” S - 62° 02’ 17” O), SW of Buenos Aires province, Argentina. The mean annual temperature in this area is 15° C and the average precipitation is 735 mm (1887-2012).

The experiment was initiated in 1986 and consisted in two adjacent plots with different tillage systems. The systems used in the experiment were the conventional tillage (CT), with chisel and disk harrow, and no-tillage (NT), with chemical weed control (Martinez et al., 2020). In June 2013, the plots were subdivided and one part continued with their historical tillage (CT, NT) and the other part passed to the contrary tillage system. NT passed to CT and CT passed to NT, and the new treatments are known as new conventional tillage (nCT) and new no-tillage (nNT). Historical and new treatments are described in detail in Gabbarini et al., (*in revision*).

The soil sampling was performed in 2015, at wheat emergence (December). Five plots (5 m × 5 m) away from each other by at least 150 m were randomly selected in each management condition at a 0-5 cm depth. At each plot, composite soil samples were randomly collected using a core sampler (diameter 2.2 cm) mixing 25 random cores after removing litter from the surface. The soil cores were mixed to get composite samples (ca. 500 g) in plastic bags and transported in cooled boxes to the laboratory. Sieving was done through a mesh (<2 mm) to remove stones and vegetable rest. After sieving, soil was divided into two parts and a subsample (ca. 100 g) was stored at 4°C in ziplock plastic bag.

### 2.2. Chemical and physical determinations

Soil organic carbon (SOC) was determined by dry combustion (LECO Carbon analyzer, LECO Corporation St. Joseph, MI, USA).

Aggregate size fractionation described by Neumann, et al (2013) were fractionated into five aggregate size classes (i.e., 2000-63, 250-63, 63-20, 20-2, 2-0.1 μm). Briefly, a total of 20 g dry weight was suspended in distilled water (soil/water ratio 1:5 w/v) and ultrasonicated with an energy input of 30 J mL^−1^. The ultrasonication was performed using an Ultrasonic Cleaner ultrasonicator (Testlab S.R.L., Argentina). The sand fraction (2000-250 and 250-63 μm) was separated from the other fractions by wet-sieving. The flow-through consisting of aggregate si es < 63 μm was aliquoted to four glass tubes and centrifuged at 50 g for 15 min at room temperature. To separate clay si e (< 2 μm) from silt si e (63-2 μm) aggregates, the supernatant containing the clay aggregate was decanted and collected in a 50 ml tube. The remaining pellets in the centrifugation tubes were re-suspended in distilled water and centrifuged again. The centrifugation and suspension steps were repeated seven times with decreasing centrifugation times, that is, 15, 13, 12 and 11 min, each twice, respectively. The last resuspended pellet was wet-sieved (< 20 μm), separating the coarse silt (63-20 μm) from the fine silt fractions (2–20 μm) in the flow-through. To enhance the flocculation of clay aggregate in the < 2 μm flow-through, MgCl_2_ (final concentration 3.3 mM) was added to the 3-L beaker and kept at 4°C over night. After decantation, the sedimented clay aggregate were further concentrated by centrifugation for 10 min at 2400 g. All fractions were dried at 40 °C and kept in the fridge until use.

### 2.3. DNA extraction

Soil microbial DNA was extracted and purified from soil samples (0.25 g) using the ZR Soil Microbe DNA MiniPrep™ (Zymo Research Corporation, USA) according to the manufacturer. Quantification and quality of the extracted DNA were specified using the NanoDrop ND-1000 spectrophotometer (Thermo Fischer Scientific, USA).

### 2.4. Quantitative PCR analyses

PCR quantifications of taxon-specific 16S rDNA were performed using primers and cycling conditions described in Supplementary Table 1. qPCR reactions were carried out on extracted soil DNA from different samples using Absolute qPCR SYBR green mixes (Qiagen Inc., USA) on an OneStep thermocycler (Life, USA). Standard curves for real-time PCR assays were developed by PCR amplifying the respective taxa by their specific primers following methods described in detail by Supplementary Table 1 (Fierer et al., 2005). Target copy numbers for each reaction were calculated from the standard curve and were used to ascertain the number of copies per g of soil. The relative fractional abundance for each of the groups was calculated by determining the copy numbers measured with each taxon-specific qPCR assay (Fierer et al., 2005). All qPCR reactions were run in triplicate with the DNA extracted from each soil sample.

### 2.5. Enzyme activities assays

Enzyme activity was measured according to the method of Marx et al. (2001), based on the use of fluorogenic MUB-substrates and microplates (Truong et al., 2019). The bulk soil and soil aggregates fractions were analy ed for β-cellobiohydrolase (CEL), N-acetyl-β-glucosaminidase (NAG), β-glucosidase (GLU), phosphomonoestearase (PME), and arylsulfatase (SUL) using 4-methylumbelliferone-β-d-cellobioside, 4-methylumbelliferone-N-acetyl-β-glucosaminide, 4-methylumbelliferone-β-d-glucoside, 4-methylumbelliferone-phosphate, and 4-methylumbelliferone-sulfatase as substrates, respectively. A moist sample (equivalent weight to 1 g oven-dry material) was weighed into a sterile jar and made up to 100 g with sterile deionised water. A homogenous suspension was obtained by vigorously stirring the suspension for 1 min. Aliquots of 20 μl were withdrawn and dispensed into a 96 well microplate (two analytical replicates sample^−1^ substrate^−1^). Buffer (80 μl) was added (100 mM MES buffer, pH 6.1). Finally, 100 μl of 1 mM substrate solution was added giving a final substrate concentration of 500 μM. Plates were kept for 5 min at 30 °C and then measured for 25 min at 1 min intervals. Fluorescence (Ex: 360 nm; Em: 450 nm) was measured with an automated fluorimetric plate-reader POLARstar Omega (BMG Labtech, Germany). Fluorescence was converted into an amount of MUB (4-methylumbelliferone), according to specific standards, thereby taking into account the degree of fluorescence quenching, through soil aggregates and organic matter, in each soil suspension (Truong et al., 2019).

### 2.6. Parameterized soil physiological profiles

The 96-well Oxygen Biosensor System (OBS) plates were used to physiologically analyze soils. The OBS plates were designed and manufactured according to McLamore et al. (2014). Plates were pre-filled with 40 μl per well of C solution to determine substrate-induced respiration. To assess background respiration 40 μl of sterile distilled water (SDW) were added instead. All the stock solutions and deionized water were filter-sterilized (<0.22 μl, Sartor, USA) and stored at 4°C before loading the plates. Fresh soil suspensions were prepared by mixing 1 g soil with 5 ml SDW in a 15 ml tube containing approximately 5 ml sterile iron beads for shaking and homogenization by ULTRA TURRAX Tube Drive (IKA, Germany) for 2 min. Slurry (160 μl) was pipetted to each well, resulting in a final well volume of 240 μl. Stock solution of C (300 μl L^−1^) were prepared in order to deliver a final concentration of 50 mg total C substrate L^−1^. Carbon source selected for the CLPP analysis were propionic acid, vanillic acid, and p-coumaric acid (Sigma, USA) and no carbon was added to measure basal respiration. C sources were previously selected based on Garland et al., (2012), selecting those carbon sources that produced most different CLPP between agricultural practices in a preliminary assay. The kinetic fluorescence data were reported as normalized relative fluorescence units (NRFU) by dividing the reading at each time point by the reading at 1 h after starting (Garland et al., 2003). Physiological response was parametrized as done before to discriminate soil use and managements (Frene et al., 2020). The peak fluorescent response (Fmax) was calculated for all samples as another indicator of respiratory activity since, assuming a constant diffusion rate; the minimum dissolved oxygen concentration should reflect the amount of oxygen consumption. The time to minimum threshold response (TMR) was defined as the time required for the NFRU to increase by 10%.

### 2.7. Statistical Analysis

The data were analyzed by one-way and two-way ANOVA using Fisher’s least significant differences (LSD_0.5_, P < 0.05) to determine significant differences among treatments within each fraction or differences among fractions. Relationships between treatments were visualized using a principal component analysis (PCA). Those statistical analyses were performed using INFOSTAT software (Di Rienzo et al., 2013).

Redundancy analysis (RDA) using the *Vegan* package (Oksanen et al., 2013), was used to determine which environmental factor were related to the composition of soil microbial communities represented by the abundance of microbial communities. Monte Carlo permutations tests were applied to compute statistical significance (*n* = 999). Finally, structural equation modelling (SEM, Grace et al., 2006) was used to compare the direct and indirect pathways between Tillage system, soil aggregate size fraction and SOC on the microbial abundances, EAs and soil physiology using *Lavaan* package (Rosseel, 2012.) and visualized with the *semPlot* package (Epskamp, 2015). The first step in SEM requires establishing an a priori model based on the known effects and relationships among the drivers of microorganisms’ abundancies (Supplementary Fig. 1). To simplify our modelling approach, we used principal component analysis (PCA) to condense our multivariate data before SEM analysis. We chose principal component that representing parameter describing the EAs within different soil aggregate size fractions (PC could explain 81.9% of total variation of soil EAs), and FMAX and TMR values within different soil aggregate fractions (PC could explain 57.7% of total variation of CLPP). Bacterial taxa abundances in different aggregate size fraction were selected as a parameter to describe the bacterial community structure (PC could explain 77.2% total variation of bacteria community structure). The fit of the model was tested by using the maximum likelihood (χ2) goodness-of-fit test with p-values and the root mean square error of approximation (RMSEA) (Schermelleh-Engel et al., 2003). Both statistical analyses were performed in R v.3.4.3 (R Core Team, 2015).

## 3. Results

### 3.1. Distribution of different aggregate size fractions and Soil Organic Carbon (SOC)

The proportions of aggregate size fractions obtained by wet-sieving method are presented at Table 1. The results showed significant differences between fractions (P < 0.001), and the interaction between fractions and tillage systems were significant too (P < 0.001, Table 2). There was not significant effect because of tillage systems (P > 0.05). The proportion of the fraction 2000-250 μm show differences between treatments according to the historical tillage management: CT and nNT were significant different from NT and nCT (*P*< 0.005). Looking for the effect of changes in tillage systems, nNT compared with CT increased 33% and 5% in 2000-250 and 250-63 μm, respectively, and decreased 11% in 63-20 μm. In comparison, nCT compared with NT decreased 15% in 2000-250 μm and increased 20% and 8% in 250-63 and 63-20 μm fractions, respectively. The distribution of the aggregates fractions showed an inverse correlation between the fraction 2000-250 μm with 250-63 and 63-20 μm fractions (*P*< 0.05 for both correlations).

**Table 1.** Aggregate size fractions distribution of soils using ultrasonic fractionation under different treatment systems. The percentage of recovery is reported for ultrasonic fractionation.

| Treatments | Aggregate size fractions |  |  |  |  | Recovery |
| --- | --- | --- | --- | --- | --- | --- |
|  | 2000-250 | 250-63 | 63-20 | 20-2 | 2-0,1 |  |
| CT | 9.82b | 37.27 a | 22.72 a | 21.64 a | 3.62 a | 95.07 |
| nCT | 18.76 a | 36.13 a | 21.43 a | 18.37 a | 4.37 a | 99.04 |
| nNT | 13.07b | 39.32 a | 20.43 a | 21.75 a | 3.89 a | 98.46 |
| NT | 22.02 a | 30.1 a | 19.74 a | 17.9 a | 3.52 a | 93.28 |
|  | C | A | B | B | D |  |

Data are means, (n = 5). Different lowercase letters indicate significant differences (P < 0.05) among tillage system within each aggregate size fraction (Fisher's LSD test) and the capital letters indicate significant differences among the five fractions.

**Table 2.** Results of two-ways ANOVA analysis with a significant value of *P* < 0.05 for effects of treatment system and aggregates soil fraction on soil biochemical activities and soil microbial structure. within each aggregate-size fraction

|  | T | A | TxA |
| --- | --- | --- | --- |
| SOC | 0.0015 | <0.0001 | n.s. |
| GLU | n.s. | <0.0001 | n.s. |
| NAG | n.s. | <0.0001 | n.s. |
| PME | n.s. | <0.0001 | n.s. |
| CEL | n.s. | <0.0001 | n.s. |
| SUL | n.s. | n.s. | n.s. |
| FMAX BR | n.s. | 0.0078 | n.s. |
| FMAX Cum | 0.0006 | <0.0001 | 0.0004 |
| FMAX Prop | 0.0008 | <0.0001 | 0.0128 |
| FMAX Vai | <0.0001 | 0.001 | 0.02 |
| Bacteria | 0.0044 | <0.0001 | 0.0073 |
| Fungi | n.s. | <0.0001 | 0.0054 |
| Archaea | 0.027 | <0.0001 | 0.0011 |
| *Actinobacteria* | 0.0096 | <0.0001 | 0.0055 |
| *Acidobacteria* | n.s. | 0.0418 | n.s. |
| *Firmicutes* | n.s. | <0.0001 | 0.03 |
| *α-proteobacteria* | n.s. | <0.0001 | 0.0012 |
| *β-proteobacteria* | 0.012 | <0.0001 | 0.0001 |

T= Treatments, A= aggregates, SOC= Soil organic carbon, CEL= β-cellobiohydrolase, NAG = N-acetyl-β-glucosaminidase, GLU= β-glucosidase, PME= phosphomonoesterase, SUL= arylsulfatase. Fmax= maximum oxygen consumption, BR= Basal respiration; Cou= Coumaric acid; Prop= Propionic acid, and Vai= Vanillic acid.

The SOC showed significant differences between tillage considering the four treatments (*P*< 0.01) and between fractions (*P*< 0.0001). There was no significant interaction between fractions and tillage treatments (Table 2). The significant differences between tillage treatments appeared between the historical tillage treatments (NT and CT) and those treatments that suffered the switch of tillage (nNT and nCT) (Figure 1). The fractions 2000-250 20-2, 2-0.1 μm presented the greatest SOC content in comparison with 250-63 and 63-20 μm fractions, among the four treatments. SOC in NT aggregate fractions of size 250-63 and 63-20 μm was statistical different from nNT treatment (*P* = 0.0048), and different for the nCT in the case of fraction 200-250 μm.

**Figure 1:**
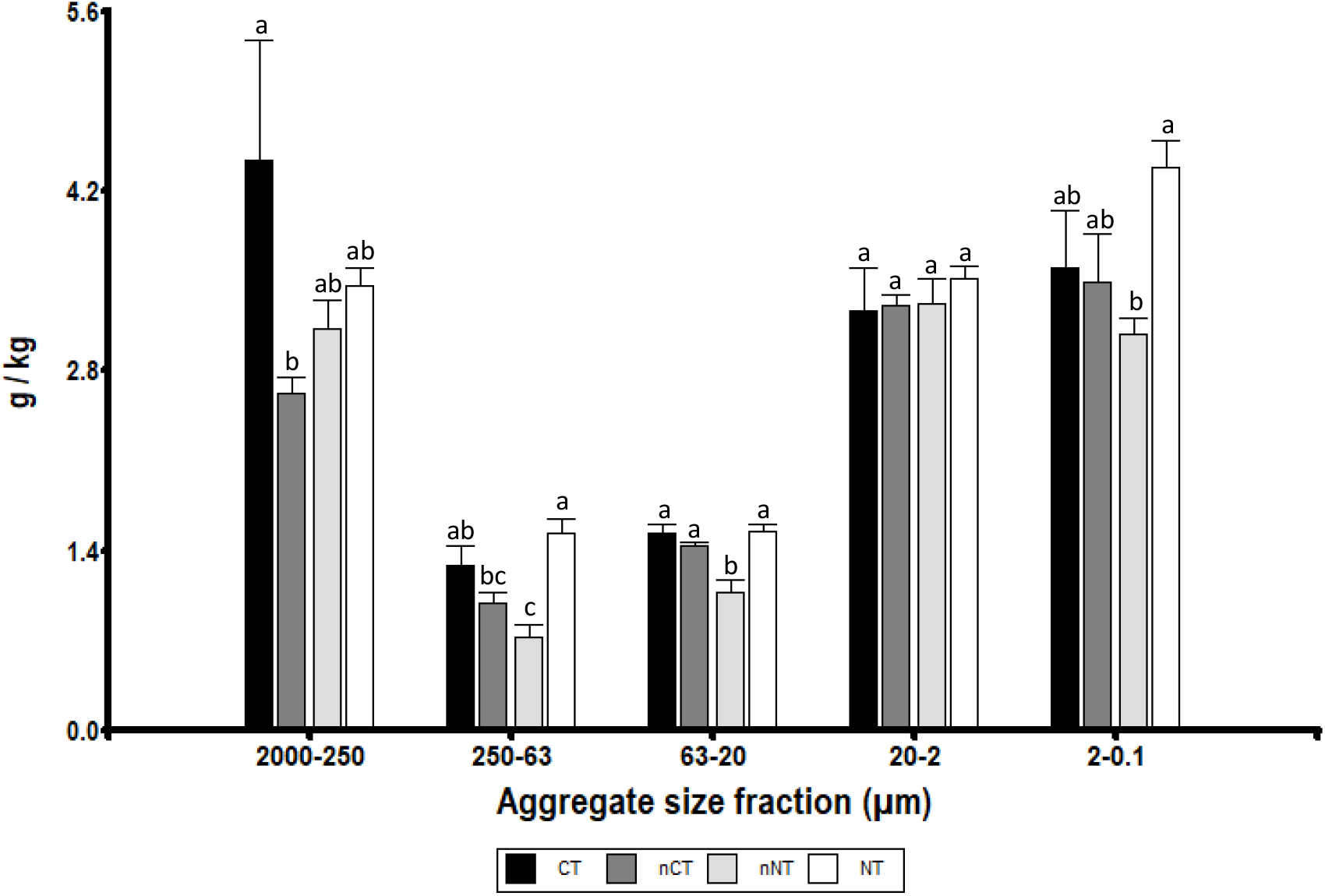
Concentrations of Soil organic C in the different aggregate size fractions under different treatments. Different lowercase letters indicate significant differences (P < 0.05) among treatments within each fraction (Fisher’s LS test).

### 3.2. Soil Enzyme Activities at the aggregate level

The 5 enzyme activities (EAs) involved in C, N, P, and S cycling were determined at the five aggregates fractions for the soils of the different tillage treatments (Supplementary Table 2). Aggregate size fraction showed significant effects on EAs (*P*< 0.05), but tillage treatments and interactions between tillage treatment and aggregate-size fraction showed not significant effects on EAs (Table 2). In general, the greatest specific activities for all enzymes were observed in aggregates fractions size 20-2 and 2-0.1 μm, following by 2000-250 fractions.; being lowest in fractions 250-63 and 63-20 μm. The impact of tillage treatments within the different aggregates size fractions showed differential results. The activities of GLU, PME and CEL were greater for CT or nCT in 2000-250, 20-2 and 2-0.1 μm fractions. NT showed the greatest activity for all the enzymes in 250-63 μm fraction. The NAG activity was greater for CT or nCT for all the fractions, with the exception of 250-63 μm. The SUL activity was greater in the NT soil samples for all the different aggregates fractions.

The PCA for five enzymatic activities of the tillage systems and aggregate size fraction (Fig. 2A) showed the CP1 and CP2 represented 81.9% and 12.9%, respectively. The soil samples were mainly separated by aggregate size fraction in three groups, the 250-63 and 63-20 μm fractions, the 2000-250 and 20-2 μm, and finally 2-0.1 μm. These groups were oriented from negative to positive values of CP1. Within 2-0.1 μm fraction, the tillage systems were grouped by actual treatment.

**Figure 2:**
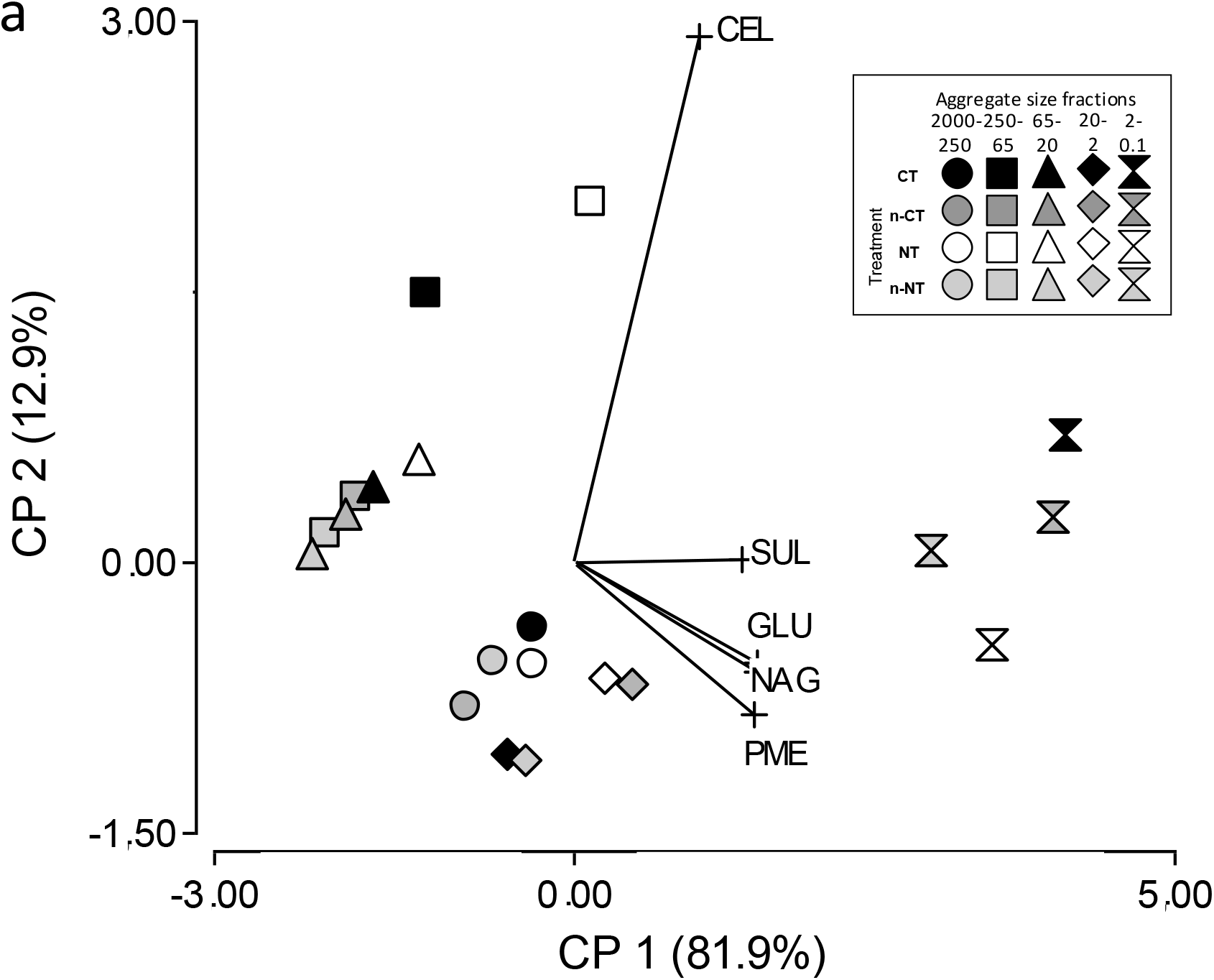
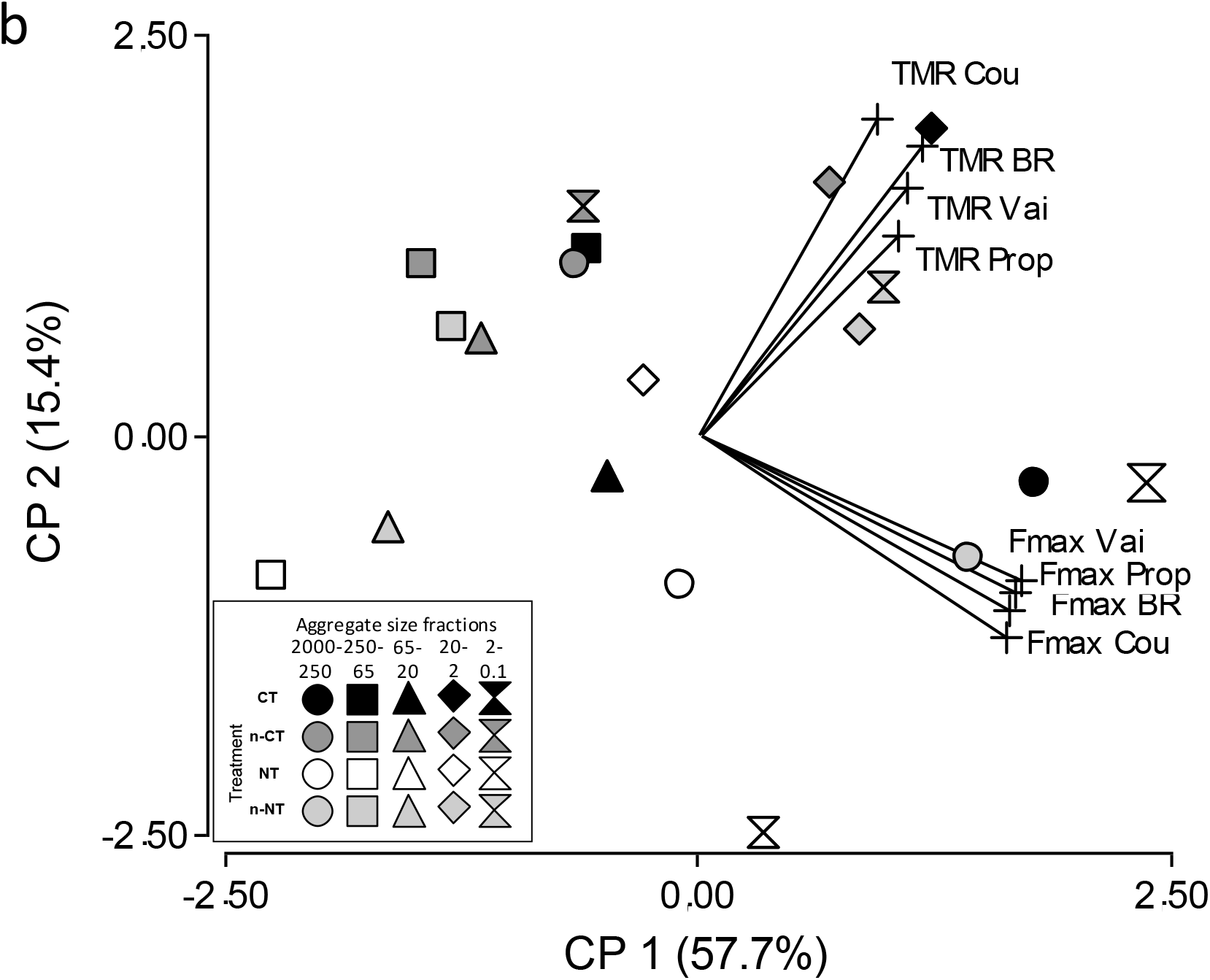
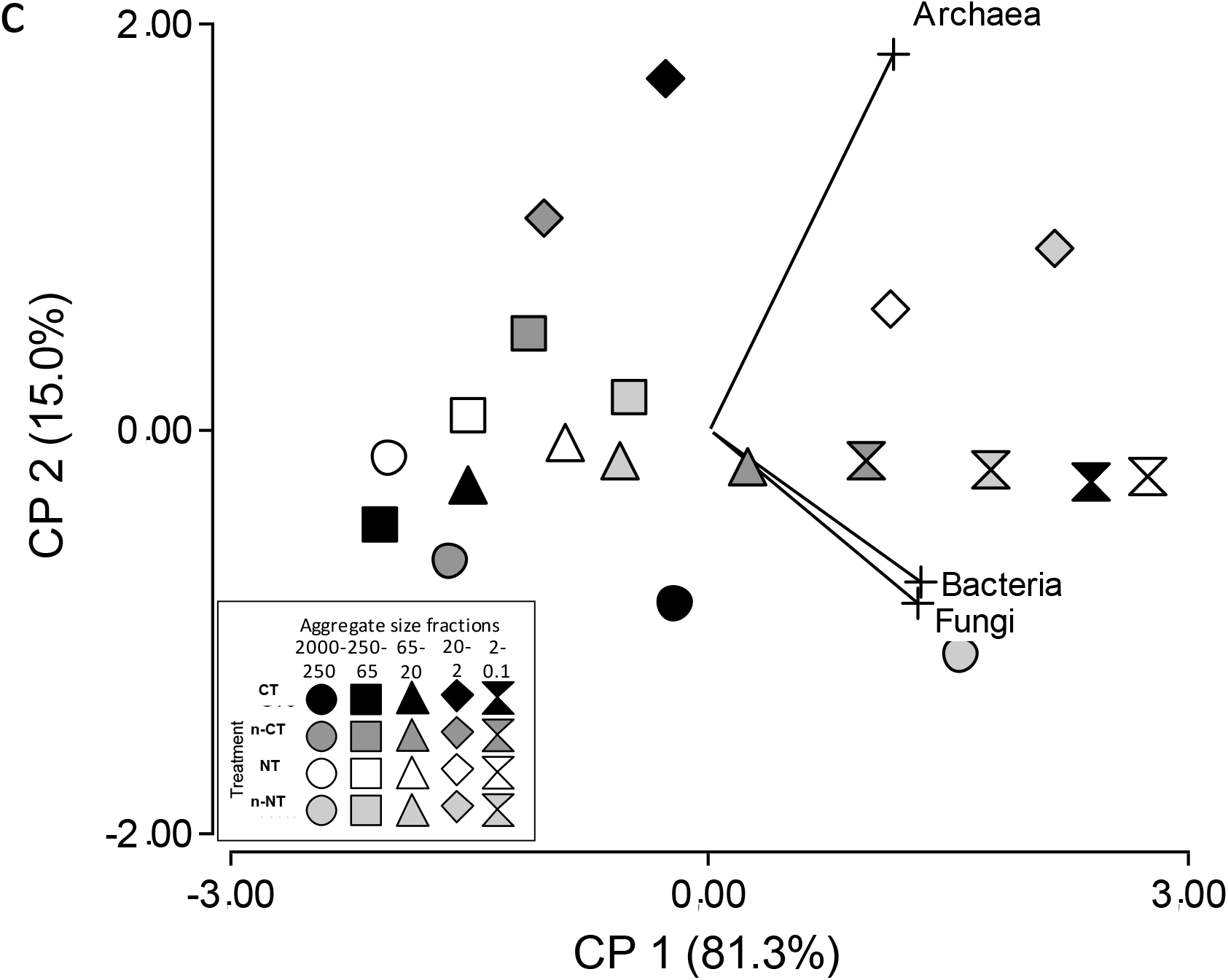
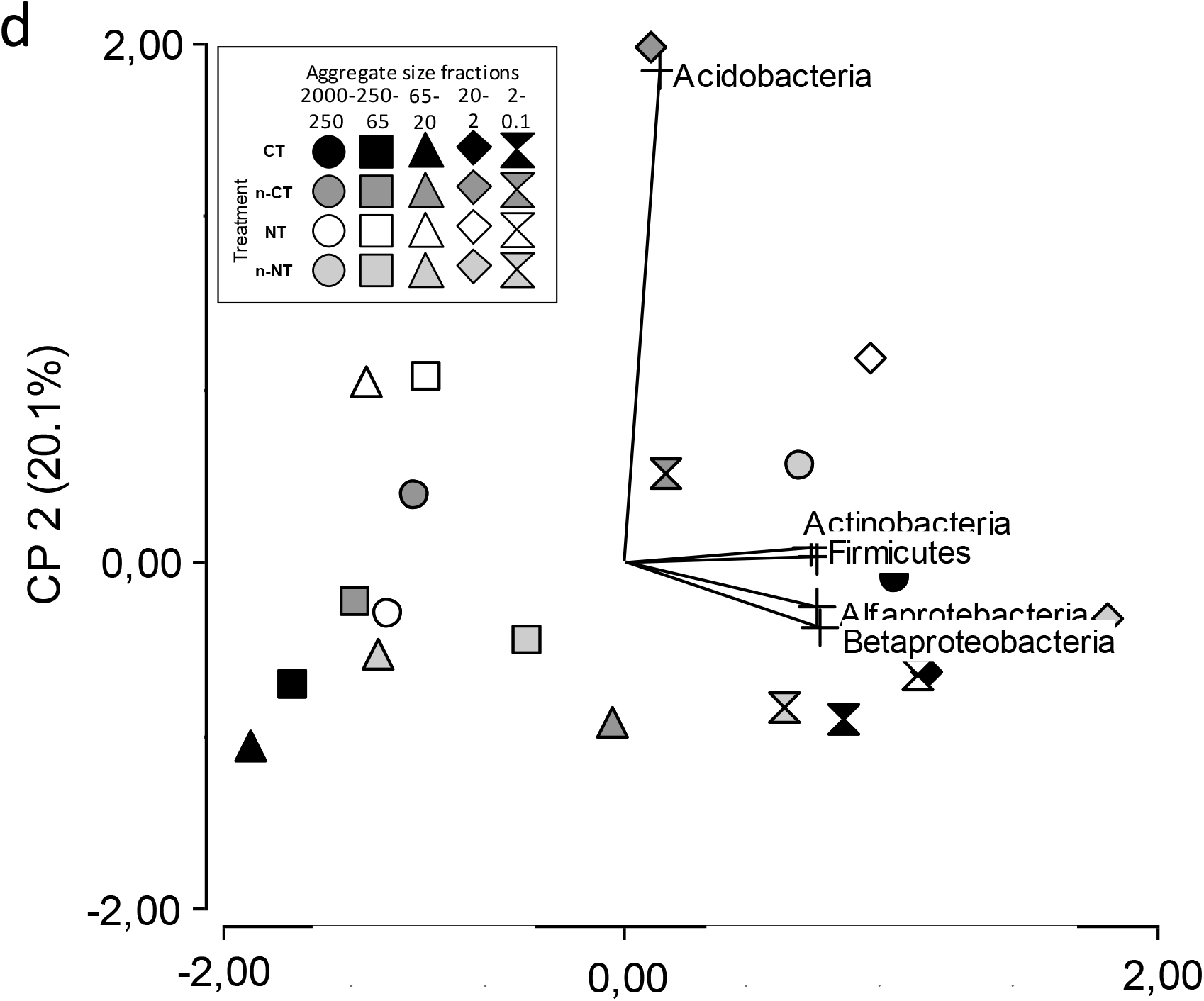
Principal Component Analysis (PCA) according to tillage system and soil aggregate size fraction for a) enzymatic profile; b) CLPP; c) soil microbial abundance; and d) soil bacterial phyla and classes abundances. Symbols: circle: 2000-250 μm; square: 250-63 μm; triangle: 63-20 μm; rhombus: 20-2 μm; Clock: 2-0.1 μm. Colors: black: CT; dark gray: nCT; light gray: nNT; white: NT.

### 3.4 Physiological profiles at the different aggregate size fractions

Parameterized physiological activity based on O_2_-consumption profiles (Frene et al, 2019) were analyzed by PCA as shown in Fig. 2B. The PC1 and PC2 explained 57.7% and 25.4% of the total variance, respectively. Substrate utilization capacity estimated through Fmax values were influenced by aggregate size fraction (*P*< 0.001) and tillage treatment (*P*< 0.001), showing also an interaction between tillage treatment and aggregate soil fraction (*P*< 0.05; Table 2). Basal respiration (no substrate addition) was influenced by tillage treatment (*P*< 0.05), but not by fraction size aggregate (Table 2). The greatest Fmax appeared at 20-2 μm aggregate size fraction (Supplementary Table 3). Considering variations within a tillage treatment, CT showed the greatest Fmax for all the fractions, excluding the activity of nNT, which was highest in 20-2 μm fraction (Supplementary Table 3).

### 3.5 Soil microbial community structure at the soil aggregate level

Soil microbial community structure was estimated by qPCR and the differences were analyzed according to the soil aggregate size fraction and tillage treatments (Table 3). Soil aggregate size fraction affected significantly general bacterial density and all the particular bacterial groups, phyla or classes, abundances (*P*< 0.05). Tillage treatments were only statistical significant for bacterial abundance (*P* = 0.0044) in terms of domains, while only phyla *Actinobacteria* and class *ß-proteobacteria* showed significant differences among tillage treatments (*P* = 0.0096 and *P* = 0.012, respectively) (Table 2).

**Table 3:** Microbial structure measured by qPCR based on in aggregate size fraction in the different tillage systems: old conventional tillage (CT), new conventional tillage (nCT), no tillage (NT), new no tillage (nNT).

| Aggregate | Treatment System | Bacteria |  | Fungi |  | Archaea |  | Acidobacteria |  | Actinobacteria |  | Firmicutes |  | Alfaproteobacteria |  | Betaproteobacteria |  |
| --- | --- | --- | --- | --- | --- | --- | --- | --- | --- | --- | --- | --- | --- | --- | --- | --- | --- |
| 2000-250 $\mu\text{m}$ | CT | 1.02E+11 | AB | 5.44E+09 | A | 1.35E+10 | AB | 5.35E+09 | A | 3.52E+10 | A | 4.35E+09 | A | 2.89E+09 | A | 4.24E+09 | A |
|  | nCT | 4.92E+10 | AB | 4.18E+09 | A | 4.40E+09 | B | 5.93E+09 | A | 1.80E+10 | A | 1.76E+09 | AB | 1.46E+09 | B | 1.91E+09 | B |
|  | nNT | 1.28E+11 | A | 1.05E+10 | A | 2.58E+10 | A | 7.07E+09 | A | 4.28E+10 | A | 3.42E+09 | AB | 2.38E+09 | AB | 3.37E+09 | AB |
|  | NT | 2.98E+10 | B | 3.24E+09 | A | 8.44E+09 | B | 3.91E+09 | A | 1.64E+10 | A | 1.02E+09 | B | 1.61E+09 | AB | 2.29E+09 | B |
| 250-63 $\mu\text{m}$ | CT | 3.23E+10 | A | 3.61E+09 | A | 3.42E+09 | A | 2.28E+09 | A | 1.34E+10 | A | 9.82E+08 | A | 1.13E+09 | A | 1.63E+09 | A |
|  | nCT | 3.37E+10 | A | 4.81E+09 | A | 2.49E+10 | A | 4.05E+09 | A | 7.32E+09 | B | 1.63E+09 | A | 1.55E+09 | A | 2.04E+09 | A |
|  | nNT | 7.03E+10 | A | 4.67E+09 | A | 2.57E+10 | A | 3.70E+09 | A | 2.37E+10 | A | 1.85E+09 | A | 1.81E+09 | A | 2.95E+09 | A |
|  | NT | 5.93E+10 | A | 1.98E+09 | A | 1.53E+10 | A | 8.30E+09 | A | 1.74E+10 | A | 1.55E+09 | A | 1.53E+09 | A | 2.14E+09 | A |
| 63-20 $\mu\text{m}$ | CT | 4.71E+10 | A | 3.84E+09 | A | 1.06E+10 | A | 1.27E+09 | A | 4.13E+09 | B | 8.81E+08 | A | 1.29E+09 | A | 1.73E+09 | A |
|  | nCT | 7.82E+10 | A | 7.54E+09 | A | 2.73E+10 | A | 2.09E+09 | A | 2.95E+10 | A | 2.62E+09 | A | 2.09E+09 | A | 2.83E+09 | A |
|  | nNT | 6.85E+10 | A | 5.27E+09 | A | 2.06E+10 | A | 3.11E+09 | A | 2.14E+10 | A | 8.42E+08 | A | 1.41E+09 | A | 2.11E+09 | A |
|  | NT | 5.86E+10 | A | 4.64E+09 | A | 1.88E+10 | A | 8.01E+09 | A | 1.23E+10 | AB | 1.57E+09 | A | 1.24E+09 | A | 1.84E+09 | A |
| 20-2 $\mu\text{m}$ | CT | 5.51E+10 | BC | 3.07E+09 | B | 5.00E+10 | AB | 4.02E+09 | A | 4.05E+10 | AB | 3.34E+09 | A | 4.10E+09 | A | 4.14E+09 | B |
|  | nCT | 3.80E+10 | C | 3.34E+09 | B | 3.39E+10 | B | 1.48E+10 | A | 2.78E+10 | B | 2.34E+09 | B | 2.30E+09 | B | 2.87E+09 | B |
|  | nNT | 1.25E+11 | A | 8.18E+09 | A | 5.93E+10 | A | 5.64E+09 | A | 5.01E+10 | A | 3.35E+09 | A | 4.79E+09 | A | 5.47E+09 | A |
|  | NT | 8.93E+10 | B | 8.14E+09 | A | 4.64E+10 | B | 9.58E+09 | A | 3.73E+10 | B | 3.33E+09 | A | 3.64E+09 | AB | 3.65E+09 | B |
| 2-0,1 $\mu\text{m}$ | CT | 1.27E+11 | A | 1.17E+10 | A | 4.53E+10 | A | 3.04E+09 | A | 3.33E+10 | AB | 3.21E+09 | A | 3.57E+09 | A | 4.12E+09 | A |
|  | nCT | 9.44E+10 | A | 8.87E+09 | A | 3.45E+10 | A | 7.16E+09 | A | 2.60E+10 | B | 2.41E+09 | B | 2.54E+09 | A | 3.43E+09 | A |
|  | nNT | 1.16E+11 | A | 1.00E+10 | A | 4.06E+10 | A | 3.01E+09 | A | 3.31E+10 | AB | 3.06E+09 | AB | 2.85E+09 | A | 4.05E+09 | A |
|  | NT | 1.31E+11 | A | 1.34E+10 | A | 5.19E+10 | A | 3.98E+09 | A | 3.92E+10 | A | 3.42E+09 | A | 3.80E+09 | A | 4.34E+09 | A |

Data are mean (n=5). Different letters indicate significant differences ( $p < 0.05$ ) between tillage systems (Fisher's LSD post hoc test).

At the aggregate size level, microbial abundances variations according to the different soil tillage treatment show a particular pattern that pointed out a different dynamic in terms of changes in microbial community structure depending on the aggregate size (Table 3). Aggregates of 2000-250 μm and 20-2 μm sizes show statistically significant differences with almost all tested microbial groups, either at the domain level (Bacteria, Fungi and Archaea; three first columns in Table 3) and almost all bacteria phyla and classes tested: *Actinobacteria, Firmicutes, α-Proteobacteria, ß-Proteobacteria* (highlighted as grey rows in Table 3). On the contrary, aggregates of sizes 250-63, 63-20 and 2-0.1 μm show no variations at the domain level and almost at none of the phyla and bacteria classes (highlighted as white rows in Table 3), except for *Actinobacteria*. In terms of Bacteria phylum, *Actinobacteria* appeared to be the most changing group because of tillage treatments showing statistically differences at all aggregates’ sizes level, except for macroaggregate 2000-250 μm (highlighted as grey column in Table 3).

The changes of the different tested microbial groups at a particular aggregate size level showed different pattern of variations. The 20-2 μm aggregates fraction show particular differences between tillage treatments for each domain group: Fungi abundances changed according to tillage management either as historically or new one after shifting treatments (darker gray and black letters); Bacteria abundances show differences for almost all tillage treatments, being historically or not (lighter gray and black letters); while Archaea show similarities according to the historic tillage management, suggesting some soil memory (darkest gray and white letters). At the bacterial phylum or class group level, *Firmicutes* show differences according to long term tillage management with intermediate response for the shifted treatments in the 2000-250 μm aggregate fraction; *Actinobacteria* and *α-proteobacteria* show similarities suggesting soil memory of the tillage management at the 20-2 μm aggregate fractions. All the other cases with some statistical difference in the ANOVA for tillage treatments at each aggregate size, showed a diverse pattern of variations or similarities between treatments, pointing out differential patterns of responses at different microbial groups (Table 3, gray rows and columns).

In another attempt to analyze the effect of the change of tillage systems, we analyzed the variation for each microbial group density detected in the samples of the new treatments nNT and nCT, in relation to their density in the samples corresponding to the historic or previous tillage management CT and NT, respectively. For domain groups (Bacteria, Fungi and Archaea), the relationship nNT/CT increased while the relationship nCT/NT decreased for all microbial communities in 20-2 μm fraction (Supplementary Figure 2a, 2b, 2c). At 2000-250 μm fractions a similar trend was found for Fungi and Archaea communities while there was no change for Bacteria (Supplementary Figure 2a, 2b, 2c).

### 3.6 Correlations

Redundancy analysis (RDA) was performed to analyze the correlations between the partial bacterial community structure estimated with the different groups quantified and the different soil enzymes, SOC and soil aggregates profiles, using all samples from different tillage treatments as data source. In the case of the RDA for soil enzymes, the first and second RDA axes accounted respectively for 39.91 and 11.22% of the total variation between partial microbial community structure and enzymatic activities (Supplementary Figure 3a). The partial microbial community structure showed a significant correlation with GLU enzyme activity (*F* = 9.04, *P =* 0.001) and CEL enzyme activity (*F* = 4.43, *P =* 0.012). Meanwhile, SUL, PME, and NAG showed no significant correlation. The RDA for SOC and soil aggregates fraction proportions against partial microbial community composition accounted for 29.46 and 11.58%, of the total variation within partial microbial community structure with SOC and soil aggregates fractions (Supplementary Fig. 3b). The partial microbial community composition was significantly correlated with aggregates soil proportions (*F* = 20.12, *P* = 0.001), but did not showed a statistical correlation with SOC.

### 3.7 Structural equation model

Structural equation modeling (SEM) approach was used to predict the direct and indirect effect of tillage treatments on the aggregates soil profile, SOC, bacteria abundance, partial bacterial community structure, EAs and CLPP (Sup Fig. 1). The fit values for the model are shown in Supplementary Table 4. Tillage treatments showed no significant direct effects on either aggregate fractions or soil organic matter, but showed direct (negative) effects on Bacteria abundance and CLPP, path coefficient - 0.11 and −1.24, respectively (Fig. 3). Soil aggregation fractions showed the most significant effects on enzymatic activities, bacteria density and soil organic matter, path coefficient −3.16, −0.66 and −1.31 respectively (Figure 3). SOC showed significant positive effects on enzymatic activities, CLPP and partial bacterial community structure, coefficient pathways 0.96, 2.03, 0.7 respectively. Finally, Bacteria density showed significant effects on enzymatic activities, CLPP and partial bacterial community structure, with coefficient pathway −1.72, −2.13, 3.23 respectively (Figure 3).

**Figure 3:**
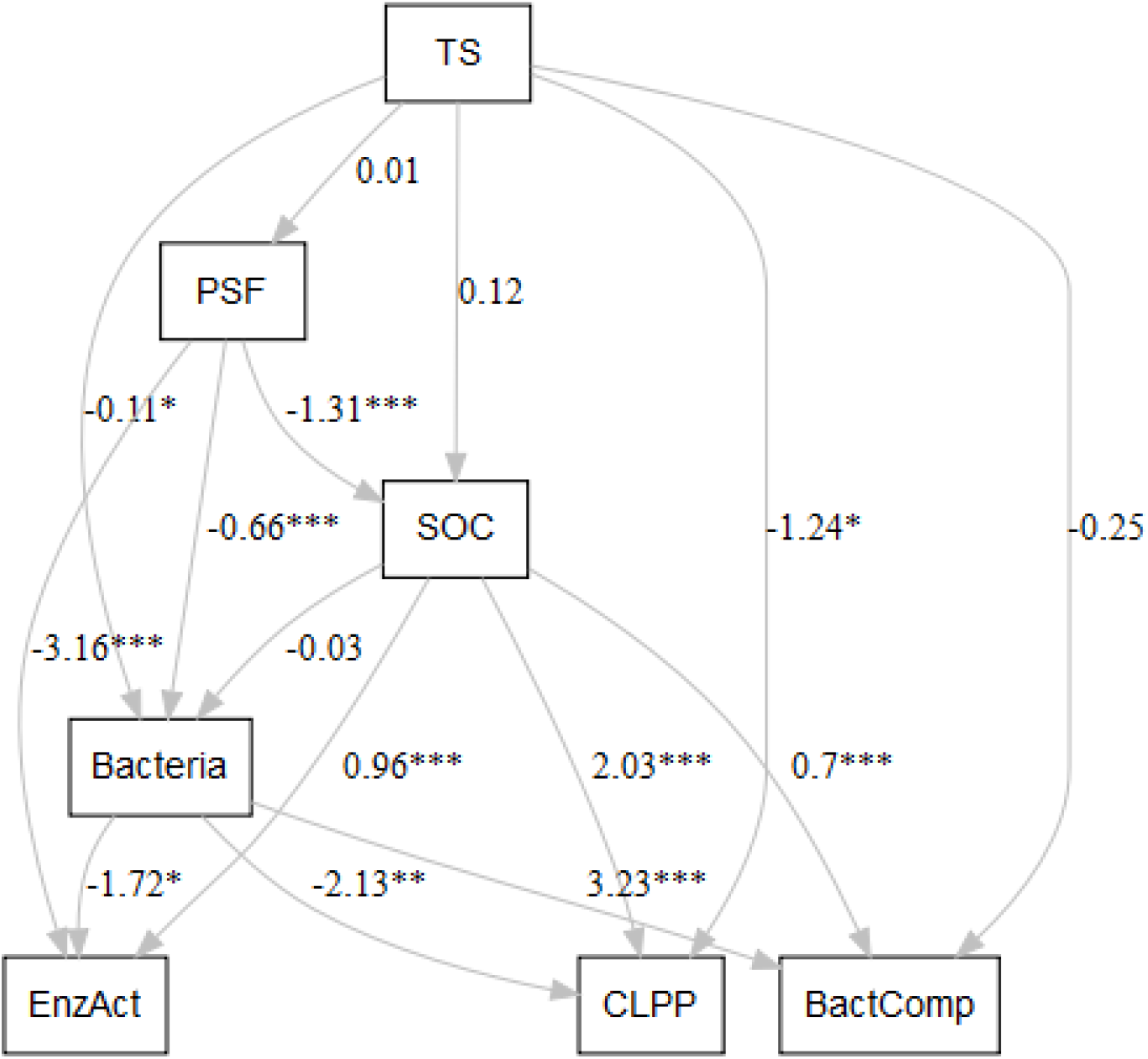
The structural equation modeling (SEM) showing the direct and indirect effects of tillage system, soil aggregates, soil organic carbon (SOC), and bacteria abundance on bacteria-related gene abundances, enzymes activity and CLPP in fractionated soils. The numbers above the arrows denote for the path coefficients. The model fits and the significance levels are presented in Supplementary Table S4. Stars denote for significance at p < 0.05, p < 0.01 and p < 0.001 probability levels (*, ** and ***, respectively)

## 4. Discussion

The switch of soil tillage managements, either from no-till to conventional tillage or from conventional tillage to no till, produce a modification in the proportion of different soil aggregate fractions. Each aggregate size fraction showed a particular microbial community structure at the high taxa level of Bacteria, Archaea and Fungi and the change of tillage managements also induced a change in this particular microbial community structure, especially in the fraction sizes 2000-250 μm and 20-2 μm. The fraction 2000-250 μm presented some changes according to tillage (nCT compared to NT, or nNT compared to CT) and lifestyle strategies (Fierer et al., 2007). When tillage is introduced in the system (nCT compared to NT) *oligotrophics* groups (*Actinobacteria* and *Acidobacteria*) increased their abundances but *copiotrophics* groups (*α*-*proteobacteria*, *ß-proteobacteia*, and *Fimicutes*) did not changed possible as a consequence of more labile C like POM. On the other hand, all phylum abundances presented increments in the nNT compared to the values in the original CT situation (Table 3). The changes of tillage systems might produce a less stable and unprotected habitat where oligotropics can quickly adapt to the conditions of the new environment in comparison with *copiotrophics* phyla (Trivedi et al., 2018). In the fraction 20-2 μm, fungal communities appear to be the community which quickly responds to changes (Frey et al., 2003; Jiang et al., 2011b) and, also, fungi are more influenced by agricultural management than soil characteristics (Wakelin et al., 2008). Bacterial and archaeal communities responded differentially to changes of tillage systems (Wessén et al., 2010).

It is worth noting the significant role of macroaggregates 2000-250 mm in the dynamics of soil structure and dynamics of microbial community in response to tillage managements, as well as the microaggregate fraction 20-2 μm. NT significantly increased the fraction 2000-250 μm in comparison with CT in a long term history of land use. Meanwhile, in the short term of 30 months, the new treatments after shift of tillage management, nCT and nNT show appeared to be in a transition state from one situation to the new one. Several authors have similarly reported greater proportion of macroaggregates in NT than CT (Helgason et al., 2010; Schutter and Dick, 2001; Zhang et al., 2016). Tillage seems to break, indirectly, macroaggregates 2000-250 μm decreasing in 15% its fraction, while increased the proportion of smaller microggregates 250-63 in 20% and 63-20 μm in 8%. This trends in the dynamics of soil aggregation had been also showed by other authors (Mikha and Rice, 2004), effects that begin to occur immediately after cultivation (Beare et al., 1994; Grandy and Robertson, 2007). On the contrary, it has been reported that NT enhanced the aggregate formation (Huang et al., 2015; Six et al., 2000), in coincidence with our results showing an increment in the amount of 2000-250 (33%) and 250-63μm (11%) in nNT compared to the original CT after just 30 months of no-till.

The change in tillage system heterogeneously modified the SOC at different aggregate levels, significantly decreasing the SOC in both new treatments nNT and nCT, suggesting a stressful transitional state from one with a long story of stabilization. Similar results have been reported in other piece of works, founding that the change from CT to NT produced a depleted of C stocks during the first 3 to 7 years (Lal, 2015b; West & Post, 2002). This change in SOC is probably linked to a change in metabolic activities of microbial communities that are also changed because of tillage systems (Phatak et al., 1999). Within the aggregates fractions profiles, CT presented greater values of SOC in 2000-250 μm (Figure 1) possible due to a 50% lesser macropore in CT than NT, resulting in a slower C diffusion into small aggregates (Smucker et al., 2007). In contrast, NT increases SOC concentration in heterogeneous way within aggregates fraction profile, being greater in middle aggregate size fractions, in coincidence with other authors observations (Dai et al., 2017). The SOC related with macroaggregates has been claimed to be related with particulate organic matter (POM), showing greater variations in comparison with 20-2 μm fraction that has been associated with mineral associated organic matter (MAOM), considered as a more stable C form which responds slowly to changes (Lavallee et al., 2020; Van Weseamal et al., 2019). It is worth noting that the differential behavior of these two aggregates fractions, 2000-250 and 20-2 μm, despite the nature of the organic matter C, is in agreement with a differential microbial community structure and change because of treatments (Table 3).

The distribution of soil EAs is often dominated by the amount and quality of organic substances as well as by various physical and chemical protection mechanisms (Allison & Jastrow, 2006). In our work, EAs presented a heterogeneously distribution with aggregate size fraction of soil (Figure 2a), following the EAs activities with the level of SOC in the fraction, as it was described in other situations (Bach and Hofmockel, 2015; Kandeler et al., 1999; Lagomarsino et al., 2012; Poll et al., 2003; Zhao et al., 2017). The EAs distribution had been explained by soil bacterial biomass, measured by qPCR (Constancias et al., 2014; Poll et al., 2003), bioactive C allocation (Lagomarsino et al., 2012), and tillage systems (Dai et al., 2017). In our case, we found that all those factors fit in a model of soil functionally interaction (Figure 3). The location of enzymes involved in C cycling, such as GLU and CEL, is directly correlated with the presence of available resources (Tiemann et al., 2015; Tiemann & Grandy, 2015) showing a close relationship with the soil microbial communities as it is suggested by the RDA result (Supplementary Figure 1b). Therefore, the EAs had been thought as closely dependent of SOC partioning among the aggregate fractions, so that SOC might serves as substrate for soil microorganisms and consequently would promote enzymes releasing into the soil (Chen et al., 2019). Following that reasoning, we found that the 2000-250 μm fractions showed the highest values of EA (except for SUL) together with highest SOC values, as is was observed in tillage soils (Li et al., 2019). CT favors soil macro-aggregates disruption, making SOC intra-aggregates available to be degraded by soil microorganisms and their extracellular enzymes (Dungait et al., 2012). On the contrary, NT would maintains and even increases macro-aggregates fraction, where SOC turnover is approximately twice as slow compared to CT (Six et al., 2000).

Microbial activity measured by O_2_-consumption CLPP reflects the size of microbial population present in soil and is strongly related with the capability of consume added substrates (Campbell et al., 2003). The relative consume of different substrates may vary among aggregates depending on microbial location and their metabolic capabilities (Jiang et al., 2011a). The microbial activity profile revealed a greater abundance of microorganisms in smaller aggregates as suggested by other authors (Hattori, 1988; Hemkemeyer et al., 2015; Lagomarsino et al., 2012; Neumann et al., 2013; Poll et al., 2003) and these results are in accordance with our qPCR quantification.

The changes on soil agricultural management, such as tillage systems or fertilization rates, alter the structure of different phylum and classes of soil microorganisms (Carbonetto et al., 2014; Finn et al., 2017; Trivedi et al., 2015). Also, soil bacterial abundances present a variation within different aggregate size fractions (Blaud et al., 2014; Trivedi et al., 2017). Our results showed that microbial abundances were different within each aggregate fraction, and a greater proportion of bacteria were associated with smaller aggregates and in a lesser way with intermediate (250-20 μm) (Constancias et al., 2014; Neumann et al., 2013; Trivedi et al., 2017; Vos et al., 2013). This distribution can be a consequence of higher concentration of fresh carbon availability, as we showed with correlation analysis for C cycling related enzymes (Supplementary Figure 3a), but we have not found a direct effect of SOC over bacteria (Figure 4). According to Neumann et al. (2013), the smaller-sized fractions provide the largest surface area in soil where microbial communities can associated with mineral particles. Therefore, it has been suggested that smaller aggregates provide a protect habitat for microorganisms through pore size exclusion of predators, like protozoa (Sessitsch et al., 2001).

Finally, the Structural Equation Model (SEM) tied together this observed variation in microbial community composition with soil and enzymes variables and tested a hypothetical model linking soil biological and physicochemical parameters with EAs, microbial activity based on CLPP and bacterial community structure (Fig. 3), pointing out that soil aggregation and SOC were not directly affected by tillage system. Other studies have similarly found that soil type and physic-chemical parameters affected microbial community composition and catabolic functions more than long-term agricultural management practices (Dai et al., 2017, Guo et al., 2018, Li et al., 2019). SEM also revealed a significant effect of SOC on bacterial community structure but not over total bacterial biomass. Davinic et al., (2012) showed that the relative abundance of soil bacteria within soil aggregate fractions are driven more by shifts in chemical composition of SOM than C quantity (content). Factors other than the chemical composition of soil, such as moisture and oxygen gradients, likely influence soil bacterial distribution and diversity (Hansel et al., 2008). If the origin of soil enzymes is mostly microbial and if soil respiration is mostly based on microbial oxygen consumption, it is clear that modification of soil structure at the microaggregate level is an acceptable explanation for soil functional changes observed because of differential soil management. The fact that neither aggregates fraction profiles nor SOC were directly affected by tillage treatments, suggests that the effect on these two soil properties are due to variables that were not measured or considered in this study, as macrofauna, mesofauna, protists, plant diversity, humidity or temperature, constitutive parts of the soil complex system.

## 5. Conclusion

Soil aggregate size fractions strongly affected the biochemical activities and microbial communities. According to this, our study showed a distinctive effect of tillage treatment of soil within each soil aggregate fraction. Thus, the changes in agricultural management modify soil microbial structure and soil services as enzymatic activities as a consequence of changes at the aggregate level of different fractions between 2000-2 μm sizes.

## Acknowledgments

This work was supported by grants PUNQ EXPTE 1411/15 and 1306/19 and project PICT 2803/17 of the Argentinean National Agency for Scientific and Technological Promotion (ANPCyT). JPF had a CONICET PhD fellowship during the period of this work sampling and soil analysis. LG and LGW are members of the Argentinean National Council for Scientific and Technical Research (CONICET). The authors are especially grateful to Christian Kleine from Hogar Funke, who took care of the long term experiment and the shift of tillage managements, and Dr. J. Galantini, who help with SOC analysis.

---

**Supplementary Figure 1:**
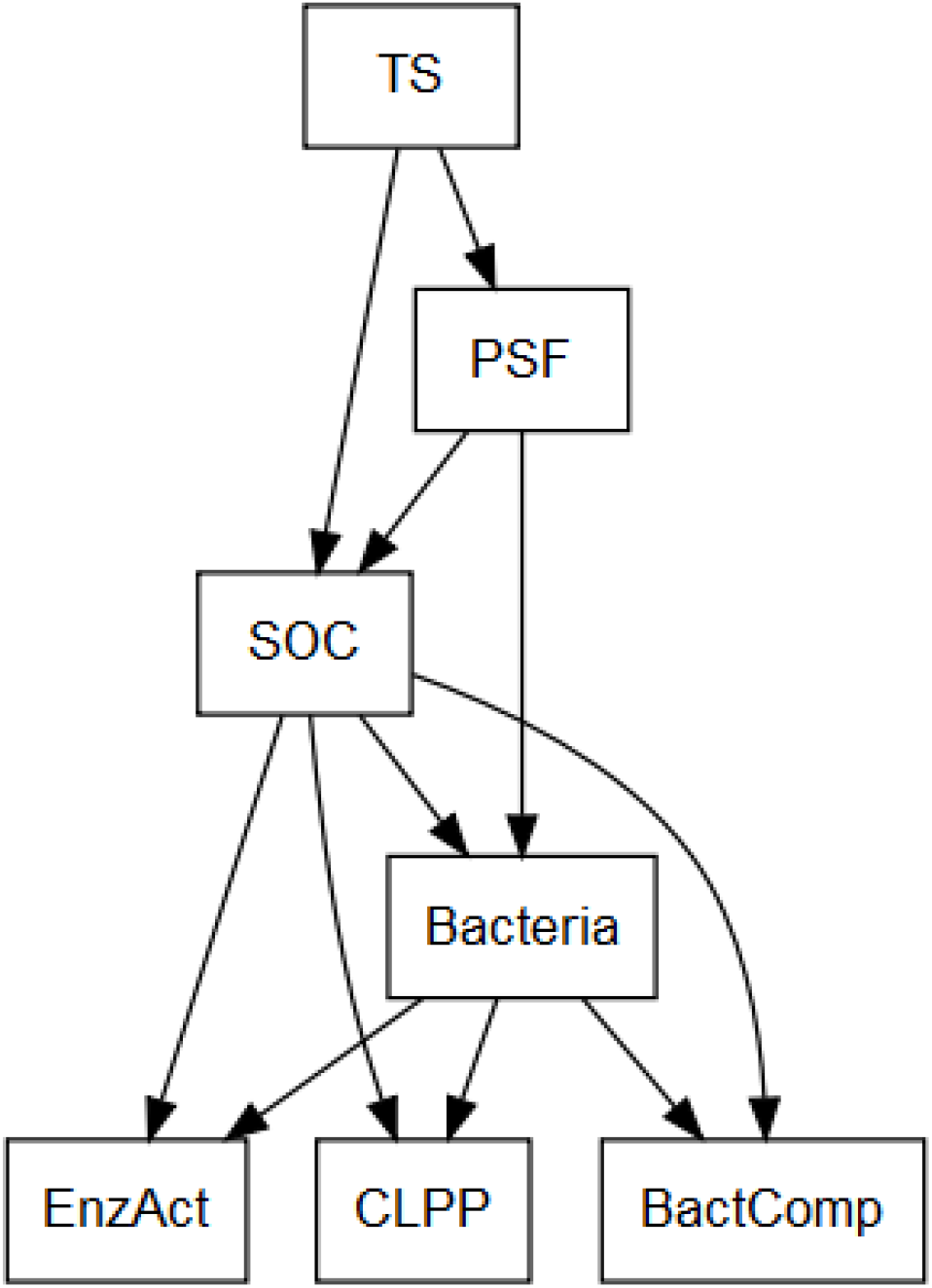
Hypothesized relationship between tillage system and the mechanistic drivers of EnzAct, BactComp, and CLPP accrual as affected by SOC and Bacteria.EnzAct = Enzymes activities; BactComp= Bacterial community structure, CLPP= microbial activity; SOC= Soil organic carbon.

**Supplemental Figure 2:**
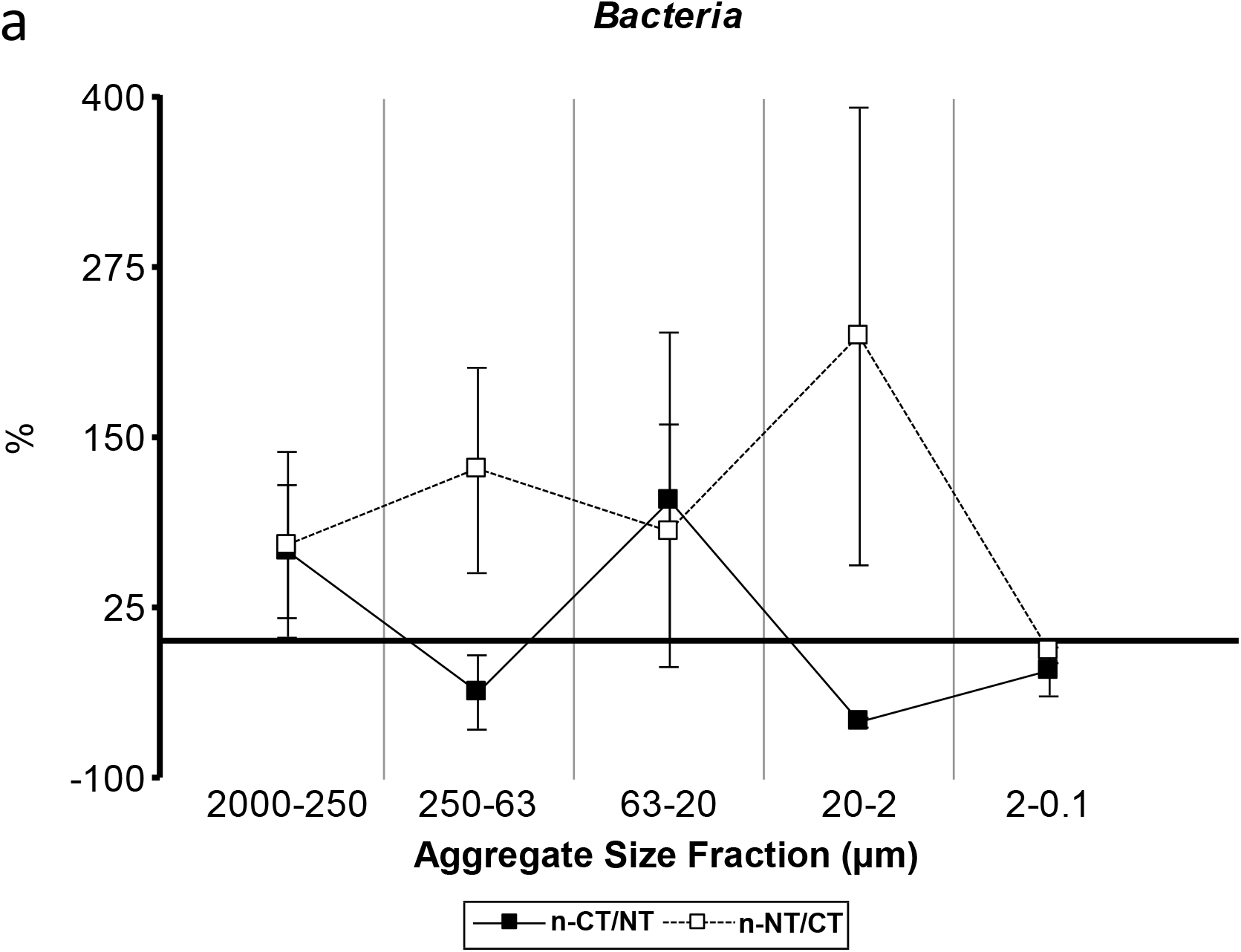
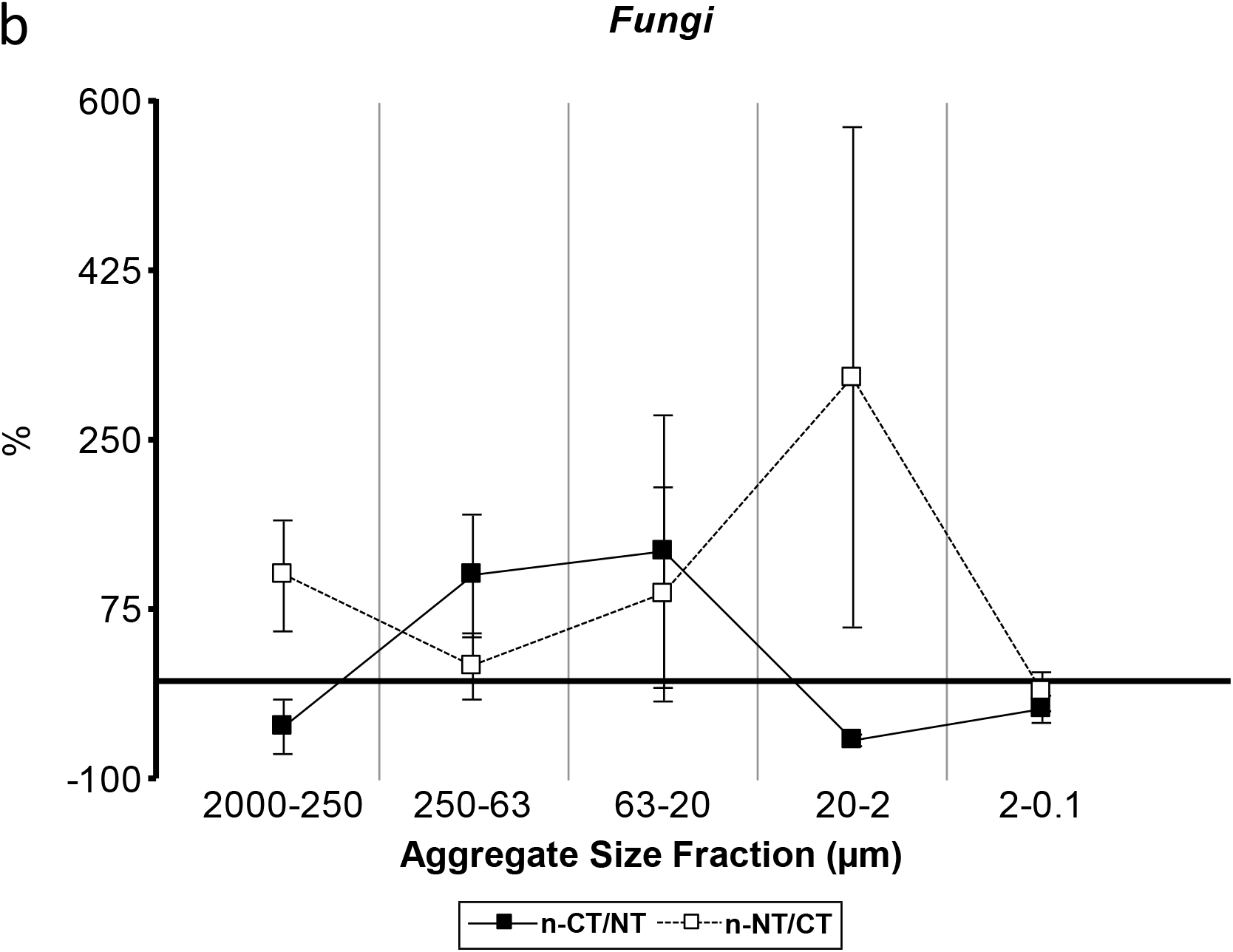
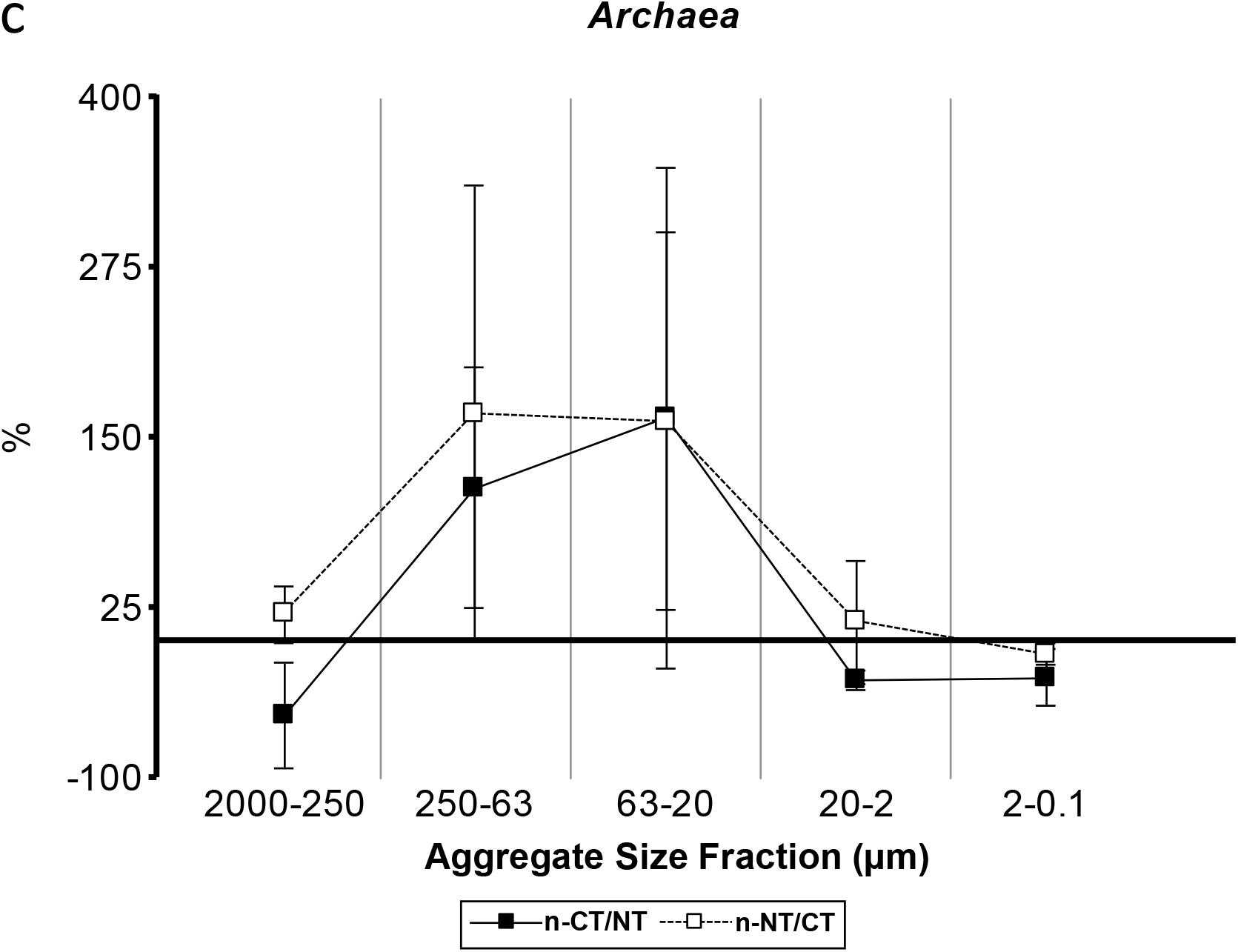
Graphics of relationships between the new agricultural management and historical tillage system predecessors for: a) Bacteria; b) fungi; c) Achaea. Colors: nNT/CT= white; nCT/NT = black (n = 5). (%).

**Supplementary Figure 3:**
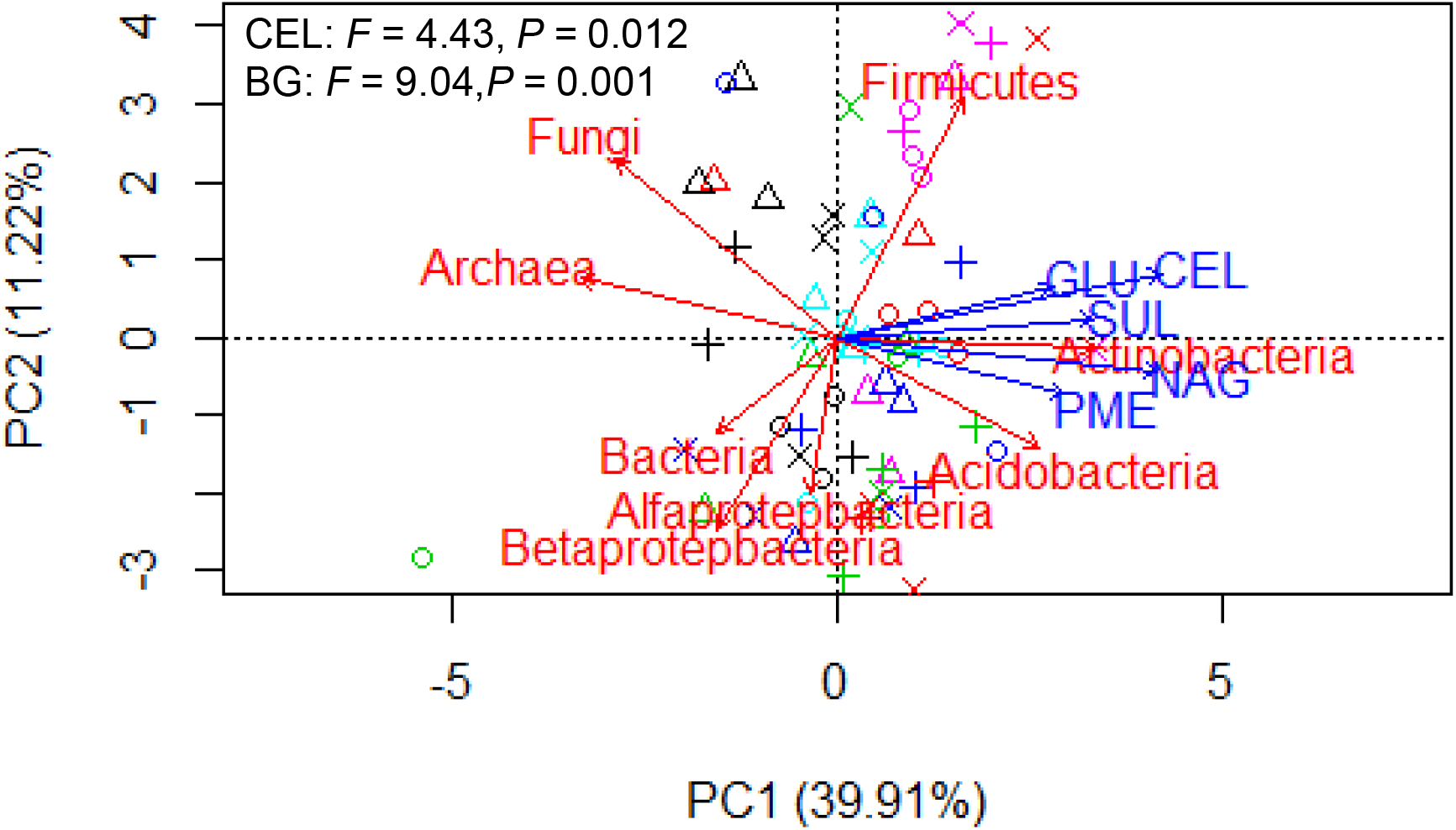
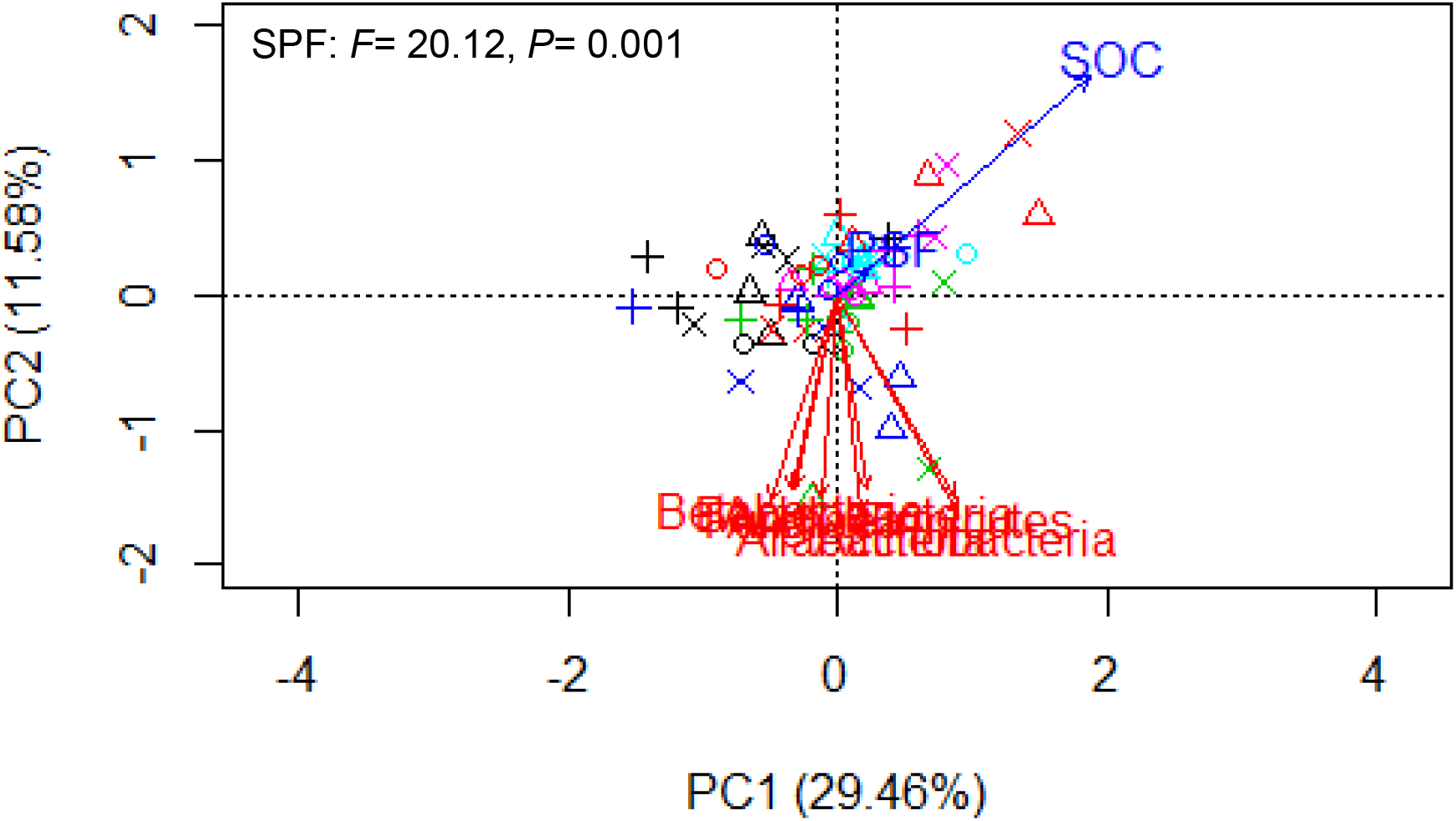
Redundancy analyses (RDA) of the correlations between (a) the correlations between microbial community composition indicated by qPCR and soil enzyme activities, (b) the correlations between microbial community composition indicated by qPCR and chemical and physical parameters. The grey arrows indicate the soil parameters and corresponding explained proportion of variability was shown in the lower right corner.

**Supplementary Table 1.**
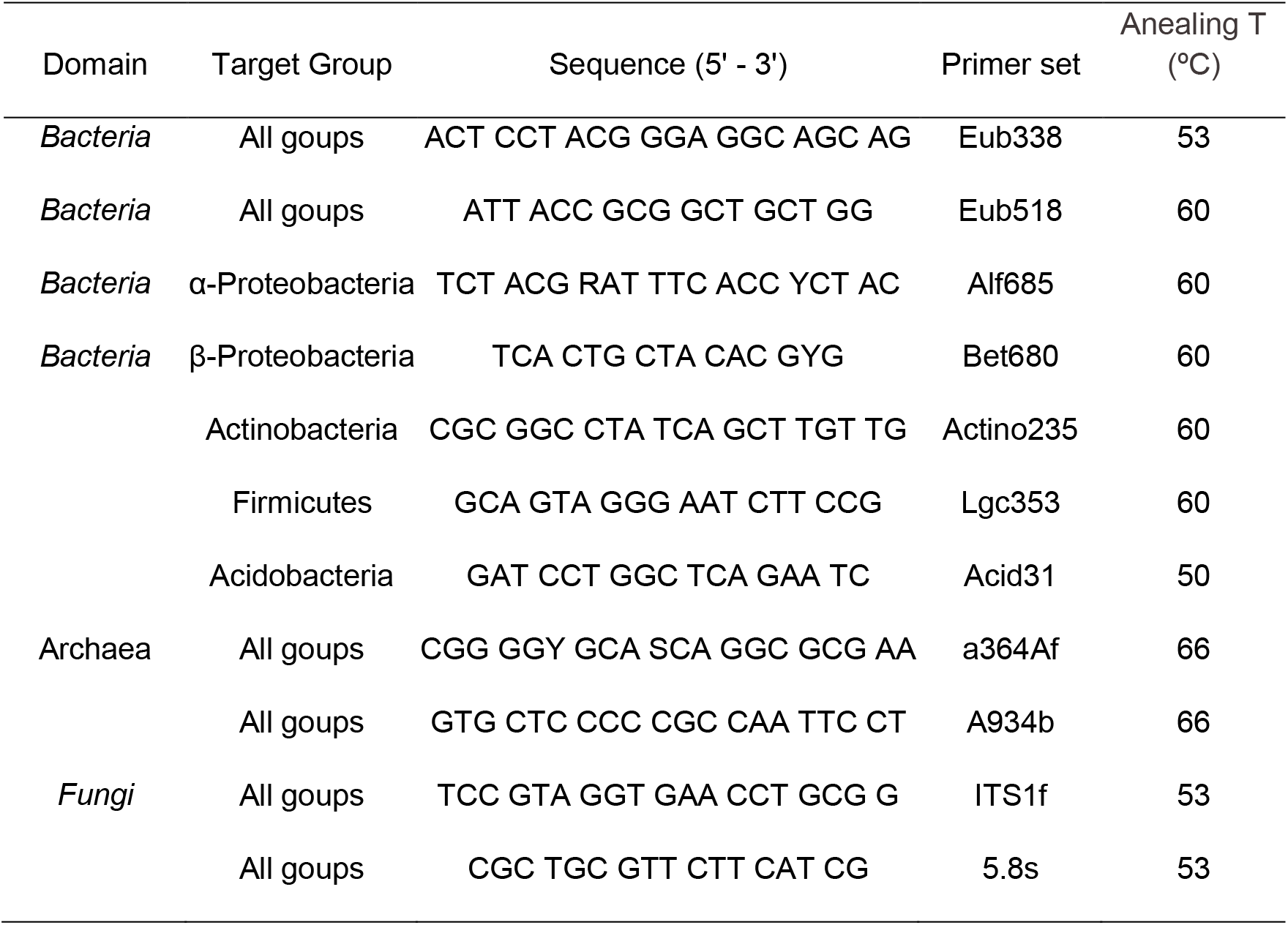
The primers used for quantitative real-time PCR.

**Supplementary table 2.**
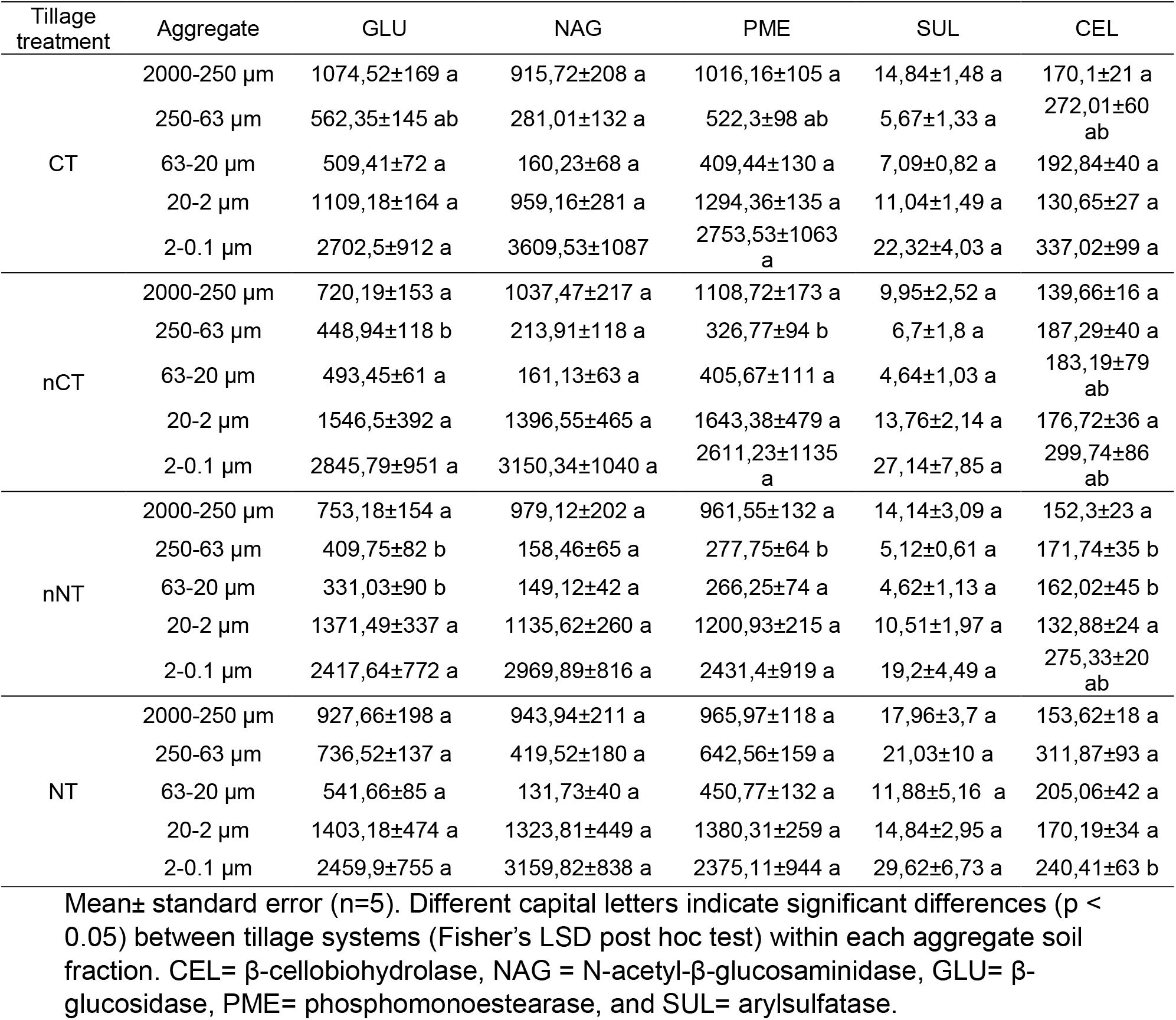
Enzyme activities (nmol MUF g^−1^h^−1^) in aggregate-size fractions in the different tillage systems: conventional tillage (CT), new conventional tillage (nCT), no tillage (NT), new no tillage (nNT).

**Supplementary Table 3.** Maximum oxygen consumption (Fmax) for each aggregate-size fraction and each different tillage treatments: conventional tillage (CT), new conventional tillage (nCT), no-tillage (NT), new no-tillage (nNT).

| Treatment System | Aggregate | Fmax BR | Fmax Cou | Fmax prop | Fmax Vai |
| --- | --- | --- | --- | --- | --- |
| CT | 2000-250 µm | 1,53±0,17 a | 1,52±0,1 a | 1,42±0,12 a | 1,57±0,15 a |
|  | 250-63 µm | 1,2±0,25 a | 1,19±0,13 a | 1,19±0,16 a | 1,34±0,18 a |
|  | 63-20 µm | 1,34±0,2 a | 1,31±0,07 a | 1,31±0,11 a | 1,32±0,09 a |
|  | 20-2 µm | 1,42±0,13 a | 1,3±0,04 a | 1,27±0,04 a | 1,35±0,08 a |
|  | 2-0,1 µm | 1,99±0,14 a | 2,02±0,23 a | 1,99±0,25 a | 2,07±0,03 a |
| nCT | 2000-250 µm | 1,33±0,07 a | 1,27±0,03 a | 1,17±0,04 a | 1,18±0,06 b |
|  | 250-63 µm | 1,2±0,08 a | 1,15±0,02 a | 1,13±0,04 a | 1,16±0,07 a |
|  | 63-20 µm | 1,25±0,06 a | 1,21±0,04 a | 1,2±0,05 a | 1,19±0,08 a |
|  | 20-2 µm | 1,38±0,08 a | 1,32±0,08 a | 1,29±0,04 a | 1,24±0,04 a |
|  | 2-0,1 µm | 1,28±0,09 a | 1,2±0,05 a | 1,25±0,06 a | 1,16±0,03 a |
| nNT | 2000-250 µm | 1,62±0,2 a | 1,54±0,11 a | 1,3±0,08 a | 1,48±0,08 ab |
|  | 250-63 µm | 1,32±0,14 a | 2±0,02 a | 1,12±0,03 a | 1,19±0,03 a |
|  | 63-20 µm | 1,34±0,13 a | 1,23±0,03 a | 1,17±0,04 a | 1,17±0,06 a |
|  | 20-2 µm | 1,48±0,13 a | 1,37±0,09 a | 1,29±0,08 a | 1,37±0,09 a |
|  | 2-0,1 µm | 1,48±0,14 | 1,34±0,1 | 1,32±0,07 a | 1,33±0,1 a |
| NT | 2000-250 µm | 1,46±0,11 a | 1,39±0,06 a | 1,31±0,07 a | 1,35±0,07 b |
|  | 250-63 µm | 1,26±0,09 a | 1,2±0,03 a | 1,16±0,04 a | 1,18±0,05 a |
|  | 63-20 µm | 1,31±0,1 a | 1,2±0,06 a | 1,15±0,03 a | 1,15±0,04 a |
|  | 20-2 µm | 1,32±0,07 a | 1,32±0,05 a | 1,27±0,07 a | 1,28±0,08 a |
|  | 2-0,1 µm | 1,5±0,09 a | 1,61±0,07 a | 1,45±0,08 a | 1,42±0,06 a |

Mean± standard error (n=5). Different letters indicate significant differences ( $p < 0.05$ ) between tillage systems (Fisher's LSD post hoc test) within each aggregate-size fraction. BR= Basal respiration; Cou= Coumaric acid; Prop= Propionic acid, and Vai= Vanillic acid.

**Supplemental Table 4.** Fit data for the structural equation modeling presented in **Fig. 4**. Abbreviations denote for: dF = degrees of freedom, χ^2^=chi-square (minimum function test statistic), RMSEA = root mean square error of approximation, CFI = comparative fit index.

| Fit<br>index | $\chi^2$ | P | dF | CFI | TLI | RMSEA | SRMR | AIC | BCC |
| --- | --- | --- | --- | --- | --- | --- | --- | --- | --- |
| Result | 0.770 | 0.857 | 3 | 1 | 1.056 | 0.000 | 0.009 | 1207.4 | 1264.7 |

